# Time course metabolomics and ^13^CO_2_ mapping establish BBX31/miP1b mediated metabolic readjustments conferring UV-B tolerance in Arabidopsis

**DOI:** 10.1101/2024.03.31.587455

**Authors:** Maneesh Lingwan, Arpita Yadav, Sourav Datta, Shyam Kumar Masakapalli

**Affiliations:** School of Biosciences and Bioengineering, Indian Institute of Technology Mandi – 175075, Himachal Pradesh, India; Department of Biological Sciences, Indian Institute of Science Education and Research (IISER) Bhopal, Bhopal Bypass Road, Bhauri, Bhopal – 462066, Madhya Pradesh, India

**Keywords:** UV-B, B-Box protein, Metabolomics, ^13^C isotope labeling, Arabidopsis

## Abstract

B-box proteins (BBXs) are transcription factors that act as signal transducers in light signaling pathways. The microprotein BBX31/miP1b is known to play a positive role in promoting photomorphogenesis and stress tolerance under UV-B. However, the BBX31-mediated metabolic reprogramming to confer UV-B tolerance in plants is not well characterised. Here, we integrate metabolomics with kinetic ^13^CO_2_ tracer-based metabolic mapping, morpho-physiological and biochemical analysis to determine the metabolic rewiring in the UV tolerant genotypes. Our results suggest that BBX31 modulates the levels of photosynthetic compounds, reduces TCA cycle intermediates and enhances GS/GOGAT metabolic intermediates and secondary metabolic pathways. ^13^CO_2_ tracing studies established BBX31 modulates phenylpropanoid and GS/GOGAT pathways to divert flux towards the accumulation of UV-B protective metabolites phenylalanine, oxoproline, glutamine, and others. Although metabolomics indicated a higher accumulation of branch chain amino acids (BCAAs) under UV-B, they had negligible 13C incorporation, indicating their biosynthesis from pre-existing intermediates or via protein degradation. Further, we demonstrate that the exogenous application of phenylalanine, identified as one of the marker metabolites, confers tolerance to plants under UV-B. This study sheds light on BBX31-mediated metabolic rewiring under UV-B, which can assist targeted enrichment of metabolites and metabolic engineering to promote UV-B tolerance in plants.

**Highlight:** BBX31/miP1b modulates the levels of photosynthetic compounds, reduces TCA cycle intermediates and enhances GS/GOGAT metabolic intermediates to confer UV-B tolerance.

## Introduction

Light is a crucial environmental factor that regulates and affects the growth and development of plants at all stages of its life cycle (Yadav *et al.,* 2020). UV radiation is an integral component of sunlight that is subdivided into UV-A (315-400 nm), UV-B (280-315 nm) and UV-C (200-280 nm) based on their wavelengths (Coohill *et al.,* 1989; Caldwell *et al.,* 1989). Elevated levels of UV-B have the potential to cause severe detrimental effects on living organisms including plants leading to loss in yield (Jansen *et al.,* 1998; Kataria *et al.,* 2014; Chuwah *et al.,* 2015). Low doses of UV-B can act as a developmental cue promoting photomorphogenic development in plants. This process is mediated by the UV-B photoreceptor UVR8 (UV RESISTANCE LOCUS 8 (Kliebenstein *et al.,* 2002; Rai *et al.,* 2020). On the other hand, high doses of UV-B can initiate stress responses in plants. Enhanced exposure to UV-B has the potential to severely damage several biomolecules including, DNA, lipids, and generate reactive oxygen species causing cell damage, necrosis and death. (Parihar *et al.,* 2015; Yoshiyama *et al.,* 2013, Lingwan *et al.,* 2024). In order to combat UV-B stress, plants adopt several strategies like enhanced leaf thickness, smaller rosettes, and increased UV-B reflective or absorptive properties by synthesizing relevant metabolites (Kusano *et al.,* 2011; Robson *et al.,* 2015;). Regardless of constant progress made, the mechanism by which fundamental components in UV-B signaling with a focus on metabolism needs to be explored in more detail.

The changes during the growth of the plant in response to UV-B light are established through various signaling pathways that are controlled through the action of several transcription factors. BBX proteins belong to a family of light-regulated transcription factors. They are zinc finger transcription factors possessing one or more B-box domains at their N-terminus and known for their actions in seedling photomorphogenesis, germination, flowering, hormone signaling, and various abiotic stresses (Khanna *et al.,* 2009; Gangappa and Botto, 2014; Vaishak *et al.,* 2019; Xu, 2019). BBX31’s role has been documented in fine-tuning photomorphogenesis by forming a feedback loop with other BBX proteins BBX30, BBX28, BBX29 and HY5 (Song *et al.,* 2020, Kushwaha *et al.,* 2022). In 2019, our study showed that BBX31 is involved in UV-B signalling and *35S:BBX31* seedlings exhibit enhanced tolerance to a high dose of UV-B (Yadav *et al.,* 2019a). However, the overall metabolic programming in any UV-stress-tolerant plants are not well characterized previously.

Different growth stages in Arabidopsis like post-germinative growth and rosette growth are potentially affected by UV-B (Boyes *et al.,* 2001). To protect themselves from the harmful effects of UV-B, plants alter their metabolism and accumulate several photo-protective metabolites. Therefore, understanding the physiological and chemical response of plants towards UV-B might guide us to identify new metabolites offering protection against harmful radiation. Plant metabolite profiling captures the set of all metabolites at a particular time. The extent of variations in the metabolite profiles due to growth stages, biotic or abiotic perturbations is obtained by undertaking a metabolomics approach in which several profiles are compared using multivariate statistical analysis (Lisec *et al.,* 2006; Celeste *et al.,* 2018; Kumari *et al.,* 2018). Previous metabolomics studies were performed in Arabidopsis, which confirmed the alteration of secondary metabolism in response to UV-B and other abiotic stress (Job *et al.,* 2022). However, metabolic response in different growth stages is not comprehensively understood in Arabidopsis. Also, the physiological, metabolic readjustments and regulation of B-Box proteins specifically under UV-B are not defined. Thus, here we aim to understand system-level metabolite readjustments of the wild type and loss and gain of function mutants of *BBX31* under UV-B.

In plants, starch and sucrose are key photosynthetic assimilates that are observed for immediate impact of carbon traffic under stress (Zeeman *et al.,* 2007). Determining plant metabolism and its metabolic rate are crucial factors to understand phenotypes under stress. Although metabolic profiling of plants can shed light on the variations in the phenotype, they are not enough to provide quantitative insights on metabolic pathways. For determining the biosynthetic routes and carbon flux of the metabolites under autotropic conditions, it is essential to feed labeled CO_2_ (Hasunuma *et al.,* 2010; Koley *et al.,* 2022). The ideal time of ^13^CO_2_ labeling is when the plant is in a dynamic steady state in which old metabolites are replaced by new metabolic intermediates at a constant rate (Schwender, 2008; Ma *et al.,* 2014). The ^13^C incorporation kinetically in metabolites and biomass components can assist in understanding plant metabolism in UV-B. In this study, we examined the ^13^C carbon traffic between metabolic networks in wild-type and BBX31 mutants under visible and UV-B.

Time course metabolomics analysis can capture UV-B-driven perturbations in the metabolic homeostasis of Arabidopsis. However, the stress response of plants may be different at different growth stages, and therefore, here we distinguish the system-level metabolite readjustments at two different growth stages in Arabidopsis. The first principal growth stage is the early leaf production stage and seedlings are 9-13 days old. The second principal growth stage was just before inflorescence when rosette leaves were fully grown (21-24 days). Understanding the comparative metabolic levels, and metabolic phenotypes in different Arabidopsis genotypes in response to UV-B can provide potential biomarkers and subsequently can help in better strategies to engineer UV-B tolerant genotypes. Thus, we extended our understanding of metabolism with wild-type and seedlings overexpressing *BBX31* under UV-B. To visualize the extent of metabolomic changes, we captured the dynamic metabolism under short to long UV-B treatment. The time course dynamic response by UV-B displayed precise and even minor metabolic shifts among genotypes. We mapped an integrated view of metabolites with differentially regulated metabolic pathways by UV-B suggesting their role in stress response. In the end, ^13^CO_2_ labeling experiments were performed to shed light on the de novo metabolic pools that are modulated under UV-B on seedling and rosette stages.

## Material and Methods

### Plant growth conditions and light treatment

For all experimental studies, *Arabidopsis thaliana* L. cv. was used, all mutants and overexpression lines used in investigations are in the background of the Arabidopsis Columbia (Col) ecotype. The mutant *bbx31* and overexpression line *35S: BBX31* have been described in our previous studies (Yadav *et al.,* 2019a). Seeds were surface sterilized using 4 % (W/V) sodium hypochlorite for 3 min and washed five times with sterile water and then inoculated on Murashige and Skoog (MS) medium containing MES (0.05%), sucrose (1%), agar (0.8%), and pH 5.78. The plates containing the seeds were subjected to stratification at 4°C for 3 days. The plates were moved to a growth chamber set at 22°C, 70% humidity, and a 16/8-hour day/night cycle for growth. Further, plants were subjected to visible and UV-B light conditions as per the requirement of experiments. For the visible light experiments, the intensity of light was set to approximately 90-110 µmol/m2/s in the Percival LED22C8 growth chamber. Light intensity was measured using the Apogee Quantum meter MQ-200. A narrow band UV-B lamp TL20W/01RS SLV (Philips) was used as the source of UV-B. It has a bandwidth ranging from 305 to 315 nm with a peak at λmax 311 nm. To measure UV-B intensity, a UV light meter UVA/UVB – 850009 (Sper scientific) was employed. The intensity of UV-B used in different experiments varied between 0.5W to 4.5W/m2, depending on the specific dosage required for each experiment.

### Chlorophyll, anthocyanin, and carotenoid estimation

For measuring chlorophyll, 100 mg fresh seedlings were crushed in dimethyl sulfoxide (DMSO) followed by centrifugation at 10000 g. Then 200 ul supernatant were taken for absorbance at 647 and 664 nm using a multi-plate reader. The measurement for total carotenoids was performed by directly measuring the absorbance of the DMSO fraction at 470 nm. The amount of chlorophyll and carotenoid present was calculated using the previously established method (Arnon *et al.,* 1967; Yadav *et al.,* 2019a). Anthocyanin measurements were performed in 6-day-old seedlings. The tissue was extracted in 1 ml of 1% HCl in methanol. The sample extracts were then incubated overnight at 4°C and centrifuged at 12000 g for 10 minutes. The absorbance measured at 530 and 657 nm respectively. The anthocyanin levels were quantified in the seedling using the formula: (A530-0.25*A657) / (tissue weight in gram) (Sakamoto *et al.,* 2017).

### DPPH assay and total phenols estimation

For measuring antioxidant response, a 2,2-diphenyl-1-picrylhydrazyl (DPPH) radical-scavenging assay was performed. 20 mg/ ml lyophilized powder was taken for extraction in chilled methanol. The sample was extracted by centrifugation at 10,000 rpm for 15 min at 4°C. A varying amount of supernatant was mixed with 100 ul DPPH (0.1 mM) solution, followed by incubation in the dark for 30 min. The absorbance was taken at 517 nm for calculating antioxidant activity. Further 50% inhibitory concentration (IC50) values were calculated. Ascorbic acid (0.01-0.1 mg/ml) was used as a positive control during the assay (Maiti *et al.,* 2014; Pant *et al.,* 2023). For measuring total phenol, 20mg lyophilized powder was extracted in acetone (70 %, 1 ml). The mixture was then centrifuged at 10,000 rpm for 15 min at 4 °C. The supernatant was taken for measuring total phenolic content using the Folin-Ciocalteu method (Kupina *et al.,* 2019). 100 µl supernatant mixed with Folin-Ciocalteu reagent (10 % v/v in water dissolved, 500 µl) and left for 5 min at room temperature. Then 400 µl sodium carbonate (5 % w/v in water) was added followed by incubation for 20 min. The total phenolics were measured at 765 nm with calibration against gallic acid (Chun *et al.,* 2003; Lingwan *et al.,* 2023a).

### Experimental design for metabolomics

The first set of experiments was conducted to capture key metabolites modulating due to UV-B in the seedling stage. 12-day-old seedlings were subjected to 24 hours of UV-B treatment. Second, to investigate the metabolite changes during the rosette growing stage, 20-day-old plants were subjected to 24 hours of UV-B treatment. Kinetic metabolomics were performed to measured central metabolic pathways kinetically under illuminated conditions of either visible or UV-B. For Kinetic metabolomics, seedlings grown in typical long-day conditions (16/8) till 21 days followed by exposure to light. Plantlets harvested at different time point (1, 2, 4, 6, 8 and 16) hours after lights treatments were subjected to metabolite analysis. In each experiment, seedlings were harvested using liquid nitrogen and crushed with a mortar and pestle. The resulting fresh weight tissue powder was then lyophilized overnight to remove all moisture and stored at −20℃ for further metabolomic and biochemical analysis.

### Metabolite extraction and GC-MS profiling

The lyophilized powder of Arabidopsis (∼20 mg) samples was subjected to the extraction of soluble metabolites. 940 µl of extraction solvent (methanol, chloroform, and water in ratio of [3:1:1 v/v/v]) and 60 µl of ribitol (0.2 mg ml^-1^ stock in H_2_O) was added to each sample and extracted at 70°C in a thermomixer at 950 rpm for 5 minutes. The extracts were then subjected to centrifugation at 13000 g for 10 min at room temperature (Lisec *et al.,* 2006). Approximately 50 µl of the supernatant was transferred to a new Eppendorf tube and dried using a speed vacuum drier. All the dried extracts samples and standards were subjected to MeOX-TMS derivatization using pyridine, methoxamine hydrochloride, and N-Methyl-N-trimethylsilyl trifluoroacetamide (Lingwan *et al.,* 2022). The data acquisition was carried out using GC-MS (GC ALS-MS 5977B, Agilent Technologies) equipped with an HP-5MS (5% phenyl methyl siloxane) and column (30 m x 250 μm x .25 μm) (Shree *et al.,* 2019).

### Protein extraction and hydrolysis

For extraction of protein, cell lysis buffer was prepared using 10mM Tris-HCl (pH 8.0), 1mM EDTA, and 1% (w/v) SDS dissolved in chilled water. Protein was extracted from ∼20 mg of tissue was dissolved in 500 μL of cell lysis buffer. Then cells were sonicated to break the cells, followed by centrifugation at 8000 g for 30 min. at 4°C. Pellets were separated from the supernatant, and after separation, 250 μL of chilled TCA was added to the supernatant. Incubated for 6 hours at 4°C and after that, protein was precipitated after centrifugation at 13000 g for 15 mins at 4°C. The supernatants were discarded, and the pellet was washed with chilled Acetone twice. The pellets were dried under a fume hood to evaporate any acetone left before hydrolysis (Masakapalli *et al.,* 2011; Ha *et al.,* 2021). The extracted protein was acid hydrolysed with 500 µL 6M HCl in a heat block for 20 hours at 100°C. The obtained hydrolysates were then subjected to centrifugation at 13,000 g for 10 min. Total 100 µL supernatant was vacuum-dried at 40°C for 6 h in a speed-vacuum system. Then, MTBSTFA (N-Methyl-N-(t-butyldimethylsilyl trifluoroacetamide) amino acid derivatives were attained with subsequent GC-MS analysis (Shree *et al.,* 2018).

### ^13^CO_2_ labeling of plants and ^13^C data analysis

The parallel plant growth chamber prototype developed was run and optimised for ^13^CO_2_ feeding experiment in standard long day conditions (16h light/8 h dark), under visible and UV-B exposure. The experiment was set up for parallel ^13^CO_2_ labeling of Arabidopsis genotypes with boxes closely airtight to avoid dilution of natural ^12^CO_2_ **(Fig. 5A)**. The dead volume was calculated and optimized to ensure equilibration of ^13^CO_2_ in the boxes after step change. The rosette was exposed to the visible and UV-B light, temperature and humidity was maintained and recorded throughout the experiment using appropriate sensors. Samples were quenched at 0, 0.5, 1, 2, 4, 8, 16 and 24 hours under controlled 400 ppm ± 50 ppm **(Fig. 5B)**. For 13C data analysis, the intensity of the mass fragments of each metabolite was obtained by using Agilent ChemStation software. The ^13^C incorporation in the carbon backbone of metabolites was mass corrected using Isocor software (Millard *et al.,* 2012). The requisite equation for Isocor’s derivatives and metabolites was computed for soluble metabolites derivatized with TMS and amino acid fragments derivatized with TBDMS. The mass ion fragments that were adjusted to account for the abundance of stable isotopes were subsequently utilised to determine the mean fraction of 13C for each metabolite (Shree *et al.,* 2018; Masakapalli *et al.,* 2013).

### Metabolomics data analysis

The raw files obtained for each condition were examined for reproducibility and evaluated using Metalign software (Lommen and Kools, 2012) for baseline correction. The metabolite profiling and data analysis were conducted using commercial Mass Hunter and Chemstation software. The identification of peak spectra for both samples and standards were conducted by analyzing mass ion fragments (m/z), retention time (RT), and identifier ions. These spectra were then compared to the NIST 17 and Fiehn Metabolomics library. For subsequent investigation, only metabolites that had a probability score greater than 70% in the NIST and Fiehn metabolomics library were chosen (Kopka *et al.,* 2005). After normalizing with the internal standard, the relative peak abundances of metabolites were extracted and subsequently analyzed using fold changes. The metadata excel file was ultimately produced for the purpose of doing statistical analysis.

### Statistical, pathway analysis and mapping with transcriptome

A file containing metadata were subjected to multivariate statistical analysis (**Supplementary Fig. S1A)**. The normalised values of identified metabolites in Col-0 and *35S:BBX31* were subjected to fold change analysis. In parallel, differentially expressed genes in *BBX31* reported in our previous study (Yadav *et al.,* 2019a) were taken for joint pathway analysis. Genes that have a fold change value is more than 2 were considered as differentially expressed genes. Further, all the differentially expressed genes mapped through KEGG mapper and response were visualised through the Path view tool for active metabolic pathways (Luo and Brouwer, 2013). Finally, the impact of metabolic features about pathways is obtained from the KEGG database using the MetPa tool and the Joint pathway analysis tool from Metaboanalyst 5.0 (Pang *et al.,* 2021). Basic statistical analysis (ANOVA and unpaired t-test) was also performed using GraphPad Prism8.

## Results

### BBX31 activates UV-B protective machinery to mitigate oxidative stress

UV-B light induces morphological responses depending on its dose of exposure (Jenkins, 2014). To investigate the role of BBX31 in UV-B stress tolerance, we exposed seedlings to visible and intensity UV-B. Initially, we irradiated seedlings with 4.5 W/m^2^ UV-B light continuously for three days. We observed that at 15 days most of the Col-0 seedlings were bleached owing to chlorosis. At the same time, the mutants showed greater extents of chlorosis, whereas *35S:BBX31* seedlings were predominantly green **(Fig. 1A)**. *bbx31* mutant seedlings showed a rapid decrease in chlorophyll upon UV-B exposure whereas lesser degradation of chlorophyll was detected in *35S:BBX31* compared to Col-0 **(Fig. 1B)**. The anthocyanin levels were significantly elevated in the overexpression lines compared to Col-0 **(Fig. 1C).** Carotenoids also decreased under UV-B in all the genotypes but the *35S:BBX31* is able to retain more carotenoids compared to Col-0 and *bbx31* mutant seedlings. **(Fig. 1D).** 2,2-diphenyl-1-picrylhydrazyl (DPPH) analysis showed that IC50 value was lowered in *35S:BBX31* suggesting higher antioxidant activity under UV-B **(Fig. 1E)**. Total phenolics were significantly abundant in *35S:BBX31* seedlings **(Fig. 1F).** The Col-0 and *bbx31* mutant seedlings exhibit lower level of polyphenol than the overexpressor lines. The levels of total polyphenol and dehydroascorbate were enhanced in Arabidopsis plants after supplemental UV-B treatment. Dehydroascorbate and phytol level were measured through GC-MS. *35S:BBX31* lines have lower dehydroascorbate than Col-0 seedlings suggesting that *BBX31* overexpressor lines might experience less oxidative stress under UV-B **(Fig. 1G).** Chlorophyll degradation releases a large amount of phytol, which is derived from geranylgeraniol pathway. Phytol is an isoprenoid alcohol and its free form is known as building component of chloroplasts (Gutbrod *et al.,* 2019). *35S:BBX31* showed a significantly lower reduction of free phytol which suggests that BBX31 might have a role in the re-synthesis of chlorophyll or the synthesis of tocopherol and other phytol-derived fatty acids **(Fig. 1H).**

**Fig. 1.**
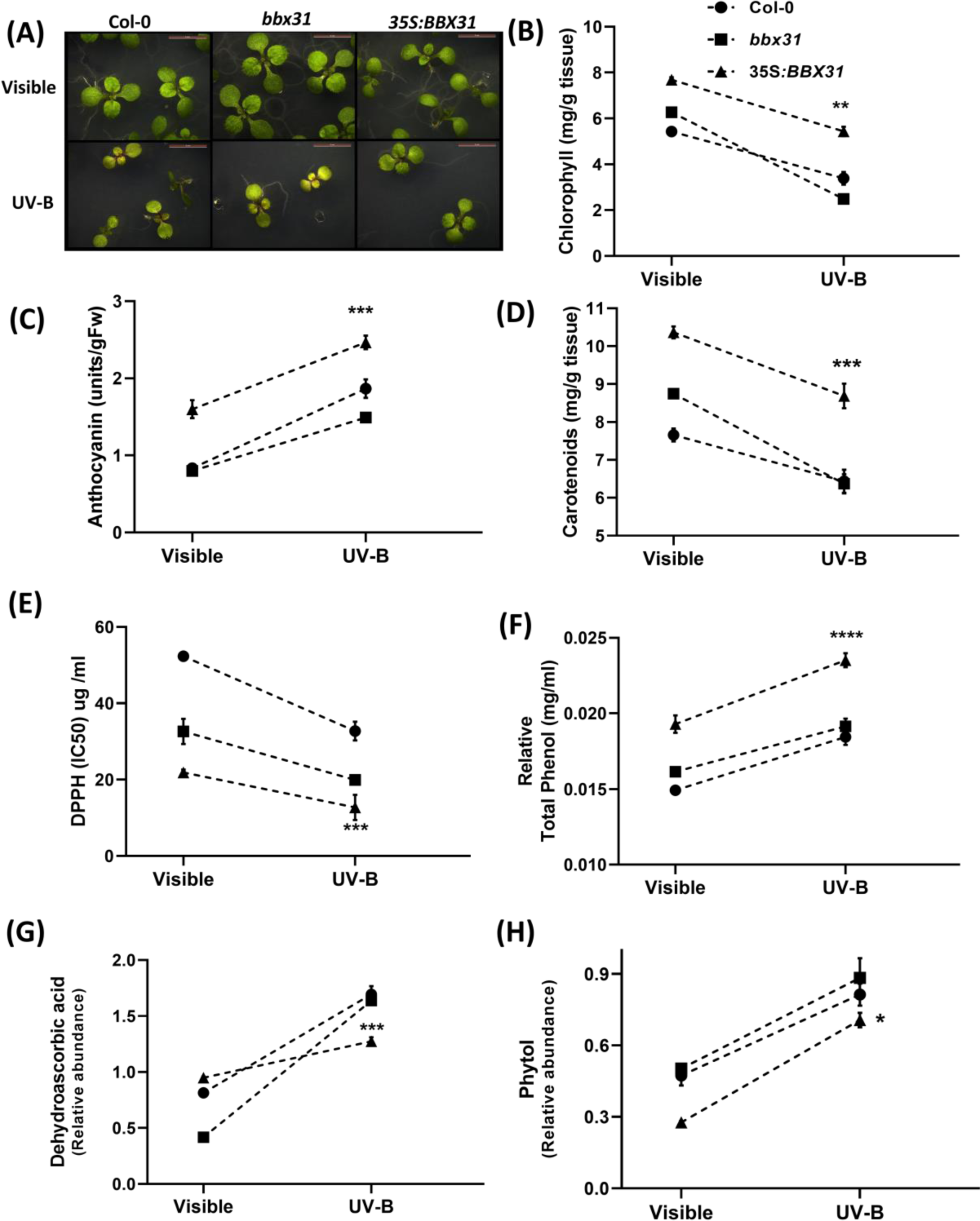
UV-B radiation lowers photosynthetic compounds and enhances antioxidant content. (A) Phenotypic analysis revealed that *35S:BBX31* lines are tolerant to UV-B radiation. Representative images of Col-0, *bbx31* and *35S:BBX31* seedlings grown in visible light (long day, 16 h/8 h) for and for UV-B treatment 12days old seedlings were given continuously for the next 3 days. (B) Chlorophyll levels lower in seedlings under UV-B. (C) Anthocyanin and (D) Carotenoid estimation showed accumulation in *35S:BBX31* seedlings in Visible and UV-B. (E) DPPH (IC50) F) Total phenolic content quantification and (G) Relative dehydroascorbic acid measurement in Col-0, *bbx31* and *35S:BBX31* seedlings indicated higher antioxidant potential in the *35S:BBX31*. (H) Relative phytol accumulation in Col-0 indicates lower biosynthesis of chlorophyll under UV-B conditions. Asterisks represent statistically significant differences determined by Student’s t-test. p-value (****P < 0.0001, ***P < 0.001, **P < 0.01, *P < 0.05 whereas ns represents non-significant (ns P ≥ 0.05).

### Metabolomics discriminates the metabolic trends in seedling and rosette stage

We captured the metabolic response of Col-0, *bbx31* mutant and *35S:BBX31* at two different growth stages. Through the untargeted approach in different light treatments, 102 metabolic features of all metabolic classes of central and secondary metabolism were identified. **(Supplementary Table S1, S2).** Multivariate statistical analysis was performed in detail to estimate the extent of variance affected by genotypes and UV-B. A volcano plot visualizes the significant difference and substantial fold change between Col-0 and *35S:BBX31*. This score plot analysis indicates succinic acid, citric acid, and asparagine significantly change during the seedling stage **(Fig. 2A)**, while fumaric, malic, and citric acid showed significant adjustments in BBX31 during the rosette stage under UV-B **(Fig. 2B).** A hierarchical heat map visualised the response of metabolites among all the genotypes **(Fig. 2C)**. UV-B treatments showed a comparable metabolic response between *bbx31* and wild-type. While comparing the response of Col-0 and *35S:BBX31,* TCA intermediates, asparagine, leucine, alanine, glucose and sucrose showed more changes under UV-B **(Supplementary** Fig. S1D, 1F, 2A). Comprehensive metabolomics using unsupervised (PCA) and supervised clustering (PLS-DA) analysis showed variations among components, and metabolite clusters shifted towards strong variations among the genotypes during the seedling stage **(Supplementary Fig. S1B, 1C).** Col-0 and *bbx31* mutants clustered together, indicating similar susceptible metabolome shifts of plants under UV-B (**Supplementary Fig. S2C, 2D).** Metabolites contributed maximum variations among the genotypes analysed through VIP score plots and Correlation analysis (**Supplementary Fig. S1E, 2B)**. The analysis depicted that amino acids and their derivatives cause maximum variations during the seedling stage in *35S:BBX31*, indicating that they may have a role in UV-B tolerance during the seedling stage **(Supplementary Fig. S1G).** In comparison, during the rosette stage, the metabolic class of sugars and organic acids contributed to maximum variations among the genotypes (**Supplementary** Fig. S2E, 3A, 3B). Further, a comparative multivariate statistical model was used to understand the metabolic response during the seedling and rosette growing stage when subjected to UV-B. A three-dimensional score plot reveals that the rosette stage has more variations among the genotypes **(Fig. 3A).** Metabolic profiles scattered in the loadings plot are succinic acid, fumaric acid, citric acid, and malic acid implying that TCA metabolites are affected between both stages **(Fig. 3B).** Further, the changes in metabolites were mapped in the metabolic network to understand pathway responses during the seedling and rosette growing stages.

**Fig. 2.**
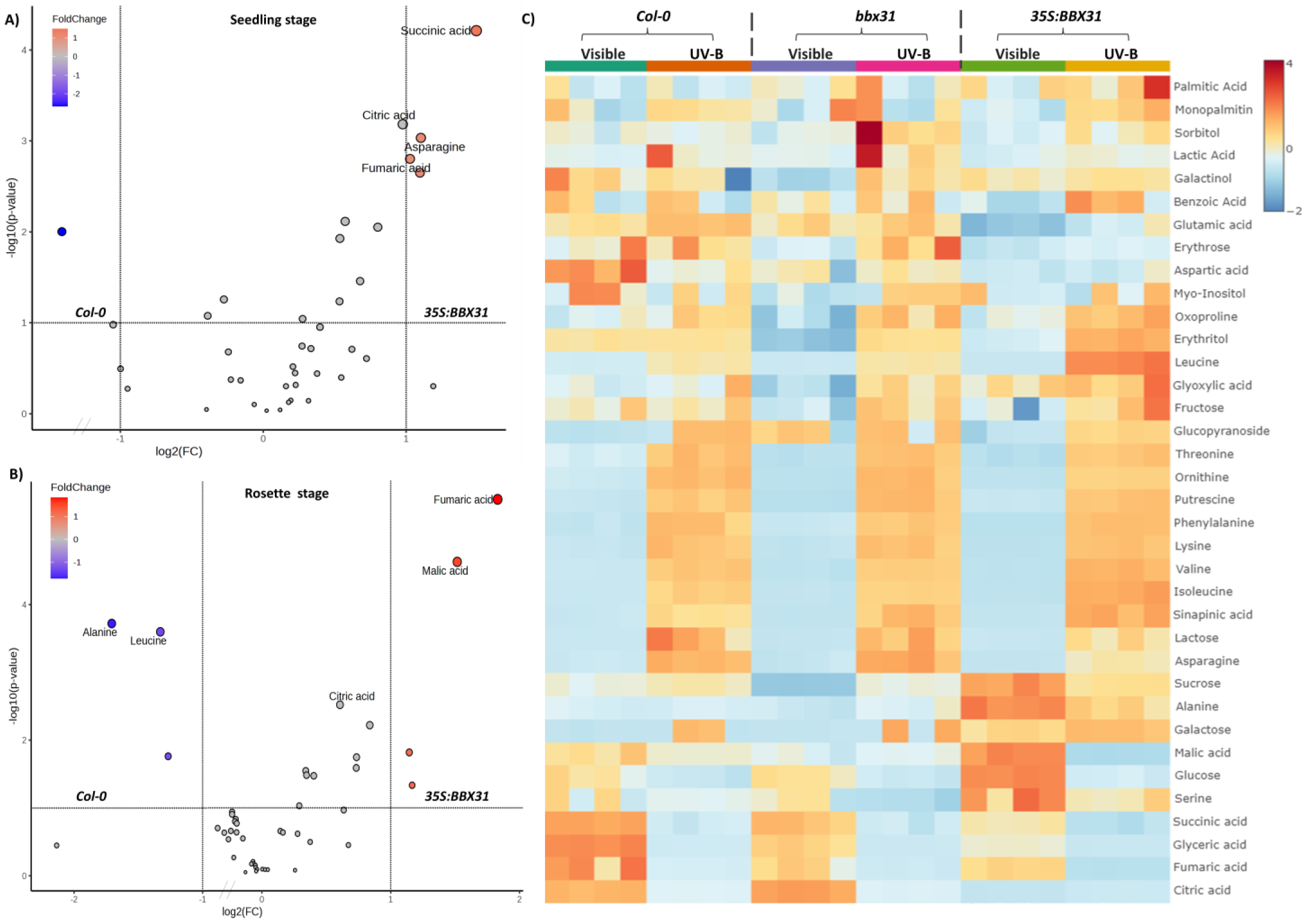
Metabolic variations in Arabidopsis Col-0, *bbx31* and *35S:BBX31* during the seedling and rosette growing stage under visible and UV-B. A) A volcano plot analysis showed TCA cycle intermediates significantly readjusted in BBX31 during the seedling and B) rosette growing stage. In the fold change scale, the x-axis represents fold changes between Col-0 and *35S:BBX31* under UV-B. In the y-axis, metabolites represented in red are significantly altered in *35S:BBX31*. C) Interactive heat map visualizing the response of metabolites in Col-0, *bbx31,* and *35S:BBX31* during the rosette stage, under both visible and UV-B treatments. Metabolite variations among the genotypes under light conditions are represented as the response scale in which blue, yellow, and red colored scales visualize from low to high levels.

**Fig. 3.**
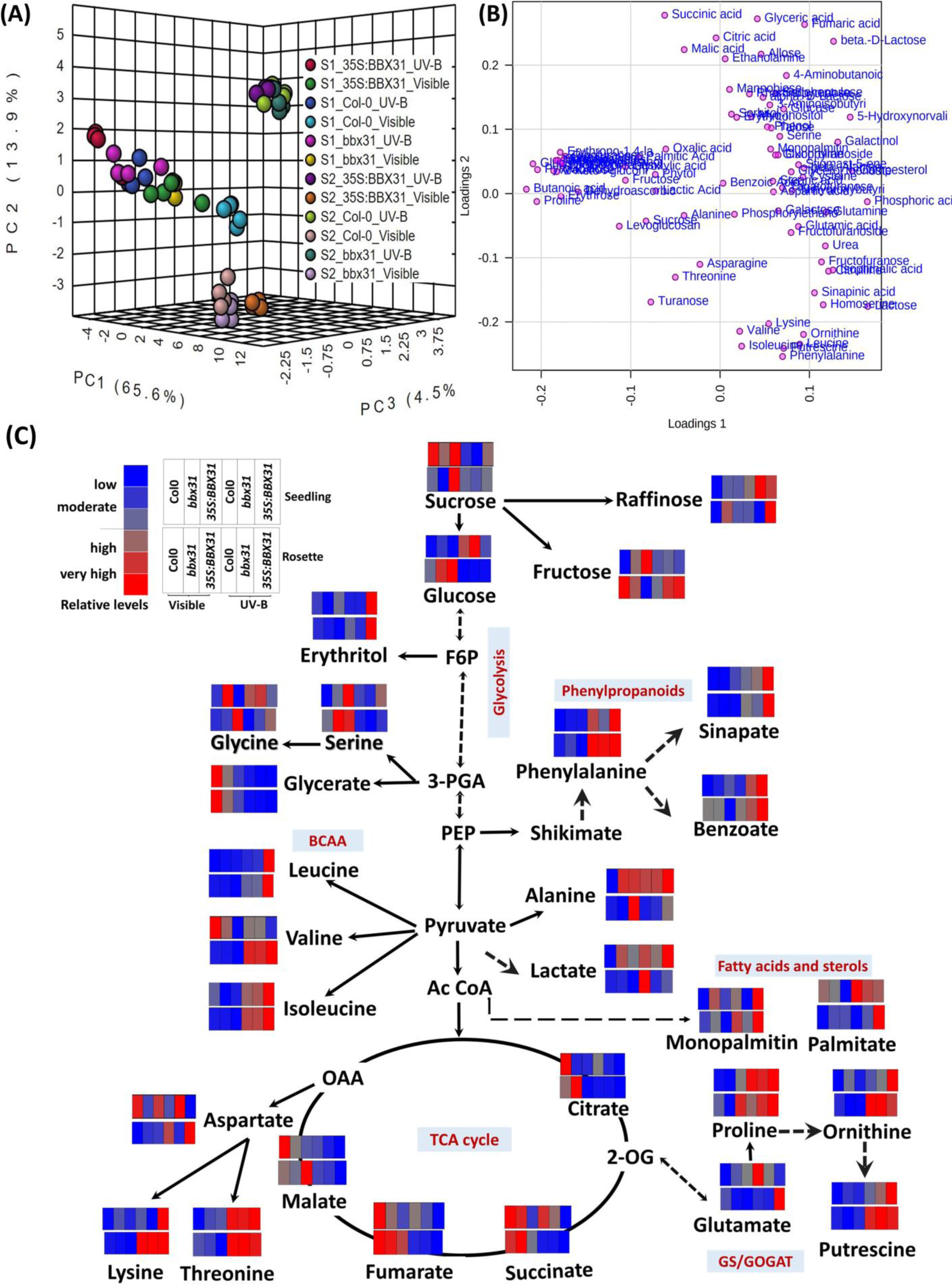
Effect of light conditions on metabolic pathways of Arabidopsis genotypes at two principal growth stages. (A) 3D score plots between the selected PCs explained variances among two stages of the Arabidopsis metabolome. The score plot reveals that the rosette stage metabolome showed more variations among the genotypes. Each circle color represents replicates, and clustering shows a set of metabolomes under certain conditions. S1 and S2 are represented as seedling and rosette stages respectively. (B) Metabolites scattered away in the loadings plot strongly influence the principal components between seedling and rosette growing stages. This represents the more scattered metabolites contributing greatest difference between the two different growth stages. For instance, metabolic profiles scattered away in the loadings plot axis are succinic acid, glyceric acid, fumaric acid, and citric acid strongly influence the principal components between seedling and rosette growing stages. C) Metabolic levels of wild-type, *bbx31* mutant and *35S:BBX31* genotypes captured and visualized in active metabolic pathways during two growing stages under visible and UV-B treatment. Relative abundance of metabolic change against UV-B is normalized with internal standard and average were means of four biological replicates. The altered pathways are shown as colored boxes, the dark red color circle has more impact on the pathway. Blue color mapped to lower impact. The upper bar represents the seedling stage, and the lower bar represents the rosette stage metabolic response. The response of metabolites in the pathway is represented in an x-axis order from 0 to 1.0. Altogether, few metabolites have similar responses in UV-B treatment irrespective of growth stage, while levels of metabolites in GS/GOGAT, sugar and fatty acid metabolism were significantly differing in *35S:BBX31* lines.

### UV-B depleted citrate pathway mediates and readjusts the amino acids

Col-0, *bbx31* and *35S:BBX31* modulate their metabolic pathways differently during UV-B stress. Intriguingly, the relative levels of citrate, succinate, fumarate, and malate were depleted in the citrate cycle (TCA) irrespective of the genotype and growing stages. However, aspartate from the oxaloacetate node was observed to be enhanced under early UV-B treatment among all the genotypes (**Fig. 3C).** The metabolic levels of amino acids derived from the aspartate node were observed to be enhanced under UV-B treatments; the increase was reflected in lysine, threonine, and isoleucine among all genotypes. However, asparagine levels were observed only to be elevated in *35S:BBX31* under UV-B. Amino acid derived from the pyruvate pool showed upregulation, and the levels of alanine, and valine elevated under UV-B, whereas leucine was significantly abundant in *35S: BBX31.* Compared with the rosette growing stage, elevated aspartate and pyruvate pool under UV-B were observed. Glycine and serine are representative amino acids of a photorespiratory cycle, synthesized in peroxisomes by amination of glyoxylate (Reumann and Weber, 2006). In both stages, UV-B affects the photorespiration however, glycine and serine levels did not show any clear trends. Altogether, the levels of metabolites in the citrate cycle were diminished irrespective of genotypes (**Supplementary Fig. S3, Supplementary Fig. S4**), while levels of metabolites from the biosynthesis node of aspartate and pyruvate were found to be significantly abundant in *35S*:*BBX31* in comparison to Col-0.

### BBX31 modulates GS/GOGAT metabolism under UV-B

The glutamine synthetase (GS) and glutamate synthase (GOGAT) pathway is altered under UV-B (**Fig. 3C).** In the seedling stage, Glutamine showed elevated levels in Col-0 but depleted in *35S:BBX31* whereas glutamic acid, ornithine, putrescine and 4-hydroxy butanoic acid were moderately increased under UV-B. Proline was dramatically increased in *35S:BBX31* under UV-B treatment but moderately enhanced under Col-0. Proline is known to be a ROS scavenger, thus a higher level of proline in the seedling stage might helping BBX31 to cope up with UV-B stress. The GS/GOGAT pathway in the rosette leaf growing stage responded differently compared to the seedling stage after UV-B treatment **(Supplementary Fig. S3, Supplementary Fig. S4**) The pool of glutamine and glutamic acid was enhanced under UV-B, whereas glutamic acid was observed to be depleted during seedling stages. The levels of ornithine, putrescine, oxoproline and citrulline was highly enhanced under UV-B.

### UV-B regulates the soluble sugar and carbohydrate metabolism

During the seedling stage, carbohydrates and sugar alcohols were observed to be less altered during early UV-B stress. The relative levels of turanose, levoglucosan, ribose, sucrose, fructose, arabinose, gluconolactone and ribofuranose fluctuated but did not show significant statistical changes in all genotypes and light conditions. However, certain sugars like glucose, erythrose and glucopyranoside were enhanced under UV-B. On the contrary, during rosette growth, the levels of sucrose and glucose were depleted, whereas fructose moderately enhanced under *35S:BBX31*. The levels of levoglucosan, fructyofuranose, turanose, glucopyranoside, galactose, tagatofuranose, erythrose and talose were enhanced but rhamnopyranose, allose, galatinol, mannobiose and frucofuranoside were depleted. The levels of sugar alcohol such as sorbitol and erythritol were increased. In summary, during the leaf growing stage carbohydrate metabolism fluctuates a lot. In contrast, the levels of the sugars are less altered during the seedling stage (**Supplementary Fig. S3, Supplementary Fig. S4**).

### BBX31 triggers wax biosynthesis and Phenylpropanoid biosynthesis under UV-B

The metabolism of fatty acid and sterols displayed higher alteration during the rosette leaf growth stage compared to the seedling stage under UV-B. The levels of stigmastene and campesterol were dramatically increased, in comparison to the levels of monopalmitin, palmitic acid and steric acid, which were moderately increased due to UV-B in *35S:BBX31*. Similar trends were observed in glycerol monostearate. *BBX31* overexpression upregulates fatty acid biosynthesis (EC:1.1.1.100), fatty acid elongation (EC:3.1.2.22), cutin and suberin biosynthesis (EC:1.11.2.3) and wax biosynthesis (EC:2.3.1.75 2.3.1.20) as determined by KEGG mapper **(Supplementary Table S3).** The Path view tool confirmed and visualised that BBX31 has a role in the upregulation of cutin, suberin and wax biosynthesis and fatty acid elongation **(Supplementary Table S4).** Integrative pathway analysis from all tools confirmed the BBX31 promotes upregulation of cutin, suberin and wax biosynthesis. Metabolite–transcript joint pathway and path view tool also revealed that *BBX31* overexpression upregulates carotenoid biosynthesis (EC:3.2.1.175), phenylpropanoid biosynthesis (EC:1.1.1.195), anthocyanin biosynthesis (EC:2.4.2.51) and tryptophan metabolism (EC:3.5.5.1). In the Shikimate-derived metabolic pathway, metabolic profiling also provides evidence that aromatic amino acid phenylalanine was elevated under UV-B stress irrespective of the genotypes but was seen to be significantly abundant in *35S:BBX31* under UV-B. Overall, through close analysis of comparative data, the metabolic response of rosette growth stages seemed to have a clearer impact on *35S:BBX31* under UV-B. However, few trends were contrasting among the genotypes, growth stages, and different light treatments suggesting two separate responses of metabolites under different times and conditions. This suggests requirement of a time course kinetic metabolomics of genotypes under UV-B. This can display minor metabolic shifts and biphasic behaviour among genotypes.

### Kinetic metabolomics exhibit distinct metabolic shifts plants overexpressing *BBX31*

Dynamic variations in the metabolic levels were assessed among genotypes under visible and UV-B. The metabolic shifts in the intermediates of glycolysis, TCA cycle, photorespiration and GS/GOAT pathway were captured kinetically for 1, 2, 4, 6, 8 and 16 hours **(Supplementary Table S5, S6).** In dynamic exposure of light, the levels of aspartate derived amino acids were relatively low in visible light. However, the levels were significantly accumulated over UV-B exposure in Col-0 and *35S:BBX31*. The levels of lysine, isoleucine and threonine were significantly up in *35S:BBX31*. The pyruvate derived amino acids valine and leucine were also kinetically modulated over time. In the *35S:BBX31*, the levels of valine and leucine were exceptionally abundant **(Fig. 4).** The dynamic metabolic levels of TCA cycle intermediates were observed to be diminished irrespective of genotypes under UV-B. The levels of citrate, succinate, fumarate and malate were observed to be lowest in *35S:BBX31*. The intermediates of GS/GOGAT and downstream metabolite (glutamine, ornithine, putrescine) levels after eight hours were enhanced and found to be distinctly higher in UV-B treated plants, implying that may have specialised role in late UV-B stress response (**Fig. 4)**. Proline levels were enhanced in early UV-B treatment in both Col-0 and *35S:BBX31* lines. In summary, we spotted that kinetic metabolomics could reveal minor alterations in metabolites shift over time.

**Fig. 4.**
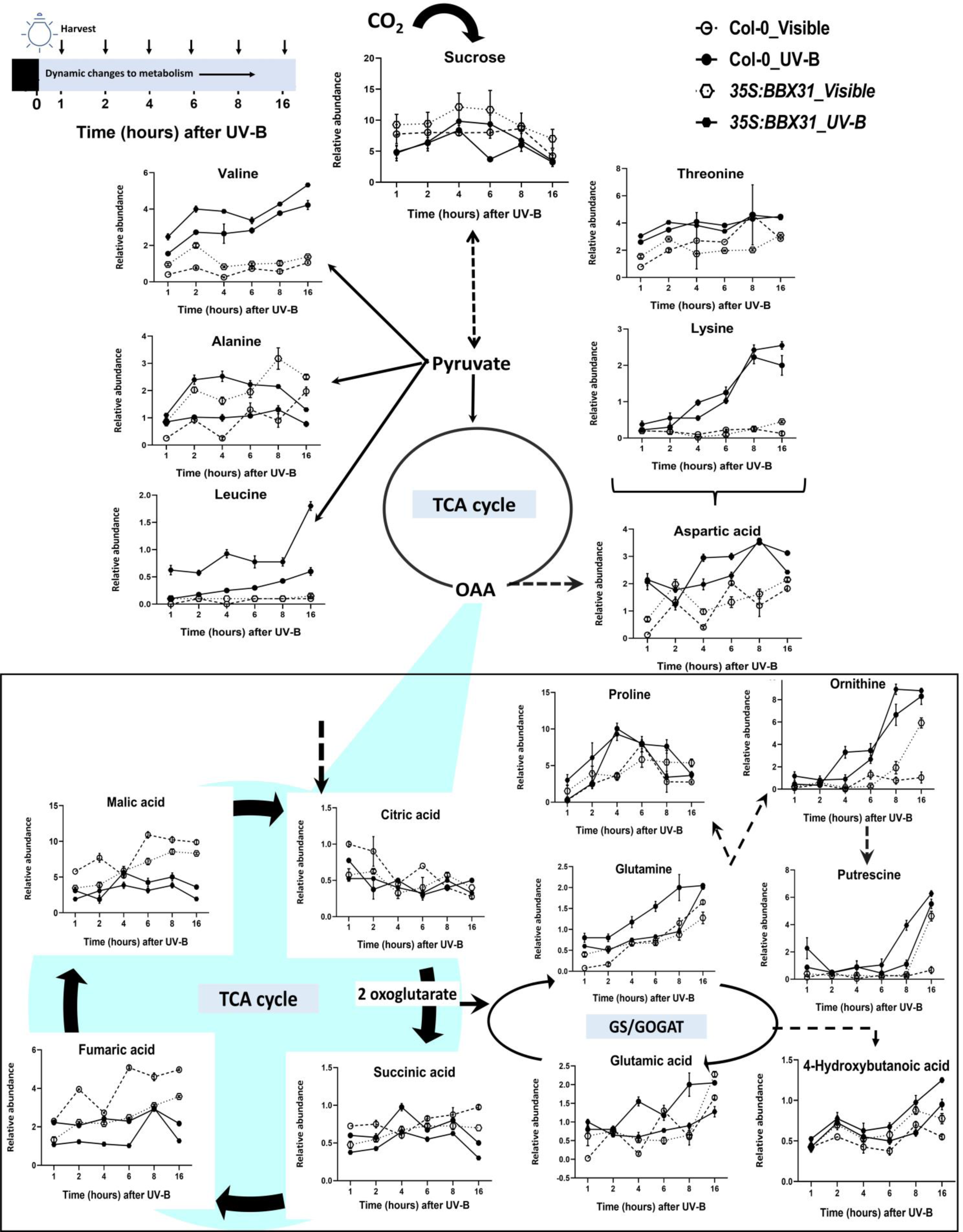
Kinetic responses of central carbon metabolism in Arabidopsis rosette stage to UV-B. The aspartate-derived amino acids were distinctively enhanced irrespective of genotypes under UV-B. The branched-chain amino acids lysine, isoleucine, and threonine accumulated in *35S:BBX31* lines under UV-B. The levels of valine and leucine were simultaneously enhanced after dynamic exposure to UV-B. The TCA intermediates were observed to be diminished in all genotypes with the lowest in *35S:BBX31*. In the GS/GOGAT pathway, proline levels enhanced in early UV-B treatment. However, the levels of glutamine, ornithine, and putrescine enhanced after eight hours of UV-B treatments, implicating their potential role in late UV-B stress response. The dotted line indicates treatments under visible light, while the dark line pattern indicates response under UV-B treatment. The y-axis signifies the relative abundance of metabolites, the circle between lines represents Col-0 response and the hexagonal signifies the *35S:BBX31* response over time. Student’s t-test and ANOVA determined statistical analyses to examine the significant differences among genotypes.

### ^13^CO_2_ tracing studies indicate presence of metabolic readjustments

Kinetic metabolomics shed light on the physiological and metabolic adaptations to visible and UV-B among the genotypes. However, whether under UV-B, the Arabidopsis genotypes fix and mobilise CO_2_ for de-novo biosynthesis of these intermediates or use pre-existing pools of TCA cycle and other intermediates was left unanswered. These queries encouraged us to quantify the metabolic phenotypes using ^13^CO_2_ analysis, which can provide insights into the carbon traffic in the GS/GOGAT metabolism and TCA cycle pathways.^13^CO_2_ feeding is the most efficient way for plants to trace the source to sink carbon flow and can define precise metabolic phenotypes (Schwender *et al.,* 2008; Allen *et al.,* 2009). The in-house parallel plant growth chamber setup was utilized for ^13^CO_2_ feeding experiment in standard long day conditions (16h light/8 h dark), under visible and UV-B exposure **(Fig. 5A, 5B)**. The incorporation of ^13^C in metabolic intermediates was analysed from the mass isotopomers of soluble metabolites and proteinogenic amino acids.^13^CO_2_ feeding experiments were conducted on Col-0 and *35S:BBX31* under visible light conditions. 13C redistribution in soluble metabolites provide detailed insights into the dynamic metabolic changes in Arabidopsis genotypes. The estimated 13C levels under visible light elevated to Alanine (50.49%), Glutamine (14.83%), Leucine (10.20%), Serine (36.42%), Sucrose (9.61%) and Valine (13.00%) in Arabidopsis Col-0 during 8 hours of ^13^CO_2_ enrichment (**Fig. 5C)**. The average 13C incorporation was observed to decrease in metabolites after 16 hours of ^13^CO_2_ feeding. Dilution of 13C hints towards utilisation of pre-existing pool of some unlabeled metabolic intermediates in the plant tissues. In parallel, the estimated 13C levels in *35S:BBX31* under visible light are highly escalated as compared to Col-0. The levels were as follows: Alanine (52.22%), Glutamine (17.75%), Leucine (13.02%), Serine (43.63%), Sucrose (15.42%) and Valine (17.49%) in *35S:BBX31* for 8 hours and similar trend was observed in 16 hours of ^13^CO_2_ enrichment. The central metabolic pathway activities were confirmed by average ^13^C incorporation in mass isotopomer of proteinogenic amino acids. The average 13C incorporation in percentage (%) among amino acids fragments were observed as Alanine (4.88%), Glycine (5.17%), Valine (3.34%), Phenylalanine (2.29%), Serine (5.74%), Threonine (1.36%), Lysine (4.29%), Tyrosine (1.68%), and Glutamine (1.96%) in Col-0. The 13C incorporation in *35S:BBX31* lines were found to be higher as average ^13^C level incorporation (%) in Alanine (14.20%), Valine (9.50%), Serine (14.745%), Glycine (11.88%), Phenylalanine (9.53%). The activity of the TCA cycle in *35S:BBX31* was confirmed by the presence of 13C incorporation in Aspartate (10.16%), Glutamine (8.1%), and Threonine (7.12%). The highest (15.1%) average ^13^C was observed in Serine (**Fig. 5D)**. The fold change analysis between Col-0 and *35S:BBX31* lines showed higher 13C incorporation in protein of *35S:BBX31* as compared to wildtype Col-0. The fold levels in Alanine (2.91), Valine (2.85), Glycine (2.30), Phenylalanine (4.16), Tyrosine (4.0), Glutamine (4.14) and Serine (2.57) were observed to be significantly higher in *35S:BBX31*.

### ^13^C labeled proteinogenic amino acids and solubles suggest disrupted photorespiratory activity under UV-B

13C incorporation in soluble metabolites provides detailed insights into the dynamic metabolic changes in Arabidopsis genotypes under UV-B. In UV-B, the average 13C levels in serine were found similar (∼10 - 11.0%) among both the genotypes up to eight hours treatment. The average 13C incorporation rises in *35S:BBX31* after 16 hours of UV-B treatment. Substantial labeling in serine implies potential photorespiratory activity in Col-0 and *35S:BBX31* under UV-B. Sucrose has average 13C enrichment of just about 2.32% and 2.73% in Col-0 and 35S:BBX31, respectively (Fig. 6A). In Col-0, the sucrose levels were found to be decreased as compared to *35S:BBX31* lines. It may because of low photosynthetic activity under UV-B. Glutamine showed average 13C enrichment of 3.89% and 9.20% in Col-0 and *35S:BBX31* respectively. The relative mass ion proportions in m/z fragments of oxoproline, putrescine, citric acid and valine were analysed for Col-0 and *35S:BBX31*. The oxoproline and citric acid fragments highlighted the higher activities in BBX31 metabolism during UV-B treatment. However, in putrescine and valine, the relative levels are found to be similar suggesting there is no de novo biosynthesis during photoperiod. The average ^13^C enrichment in mass isotopomer of proteinogenic amino acids were observed to be diminished under UV-B as compared to the visible light among genotypes. In Col-0, the average 13C incorporation in amino acid Alanine (1.85%), Glycine (1.80%), and Serine (1.91 %) were observed **(Fig. 6B)**. In *35S:BBX31* lines the average ^13^C level incorporation in Alanine (2.93%), Glycine (2.23%), Valine (1.3%), Proline (2.48%) and Serine (2.61%) were observed. The *35S:BBX31* line have moderately higher 13C incorporation as compared to Col-0 under UV-B. Interestingly in UV-B, the 13C incorporation in proline was observed only in *35S:BBX31*, whereas the label enrichment in proteinogenic proline was not observed in Col-0.

**Fig. 5.**
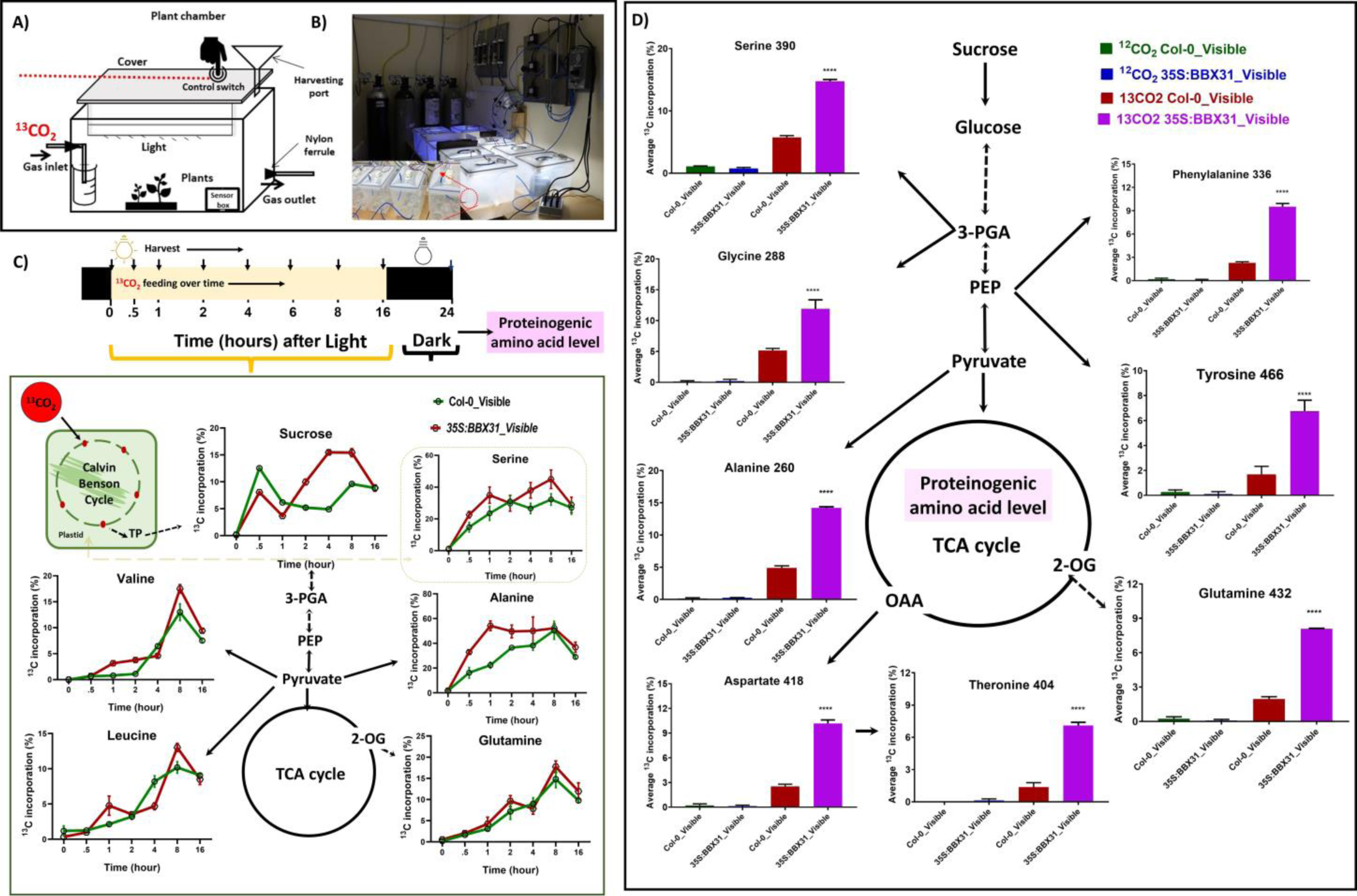
Kinetic ^13^CO_2_ mapping of Arabidopsis Col-0 and BBX31 post-light and dark phases shows metabolic readjustments with higher photosynthate by BBX31. A) The design of a box for ^13^CO_2_ feeding to plants under experimental conditions. B) The experimental setup for kinetic ^13^CO_2_ feeding of Arabidopsis seedlings. For the experiment, boxes were arranged in parallel and connected with one manifold. CO_2_, humidity and temperature sensors are calibrated, and levels are maintained during the experiments. Arabidopsis rosettes (21 d old) growing in plant growth chambers were transferred to the boxes just before the light phase started for Autotrophic phase feeding ^13^CO_2_ under the light. C) Kinetic labelling 13C observed in soluble metabolites under visible light in Arabidopsis. From each box, the rosettes were carefully quenched and harvested after giving light periods of 0, 0.5, 1, 2, 4, 8, and 16 hours. Then, samples were left at 8 hours of dark to see if the photo assimilated label into proteinogenic amino acid at 24 hours. BBX31 overexpression line compared to Col-0 showed enhanced rates of incorporation of 13C in the mass isotopomers of sugars, soluble amino acids under visible light. The 13C analysis highlighted distinct metabolic phenotypes of Col-0 and BBX31, eventually shedding light qualitatively on the activities of various metabolic pathways. D) Average 13C incorporation was higher in the fragments of proteinogenic amino acids of *35S:BBX31* than in Col-0. The higher fold levels of average 13C level incorporation in alanine, valine, glycine, threonine, tyrosine, serine, proline, aspartate, glutamine and phenylalanine were observed in BBX31 overexpression line as compared to wildtype Col-0. Statistical analyses were performed using the student’s t-test and ANOVA to examine the significant differences among genotypes.

**Fig. 6.**
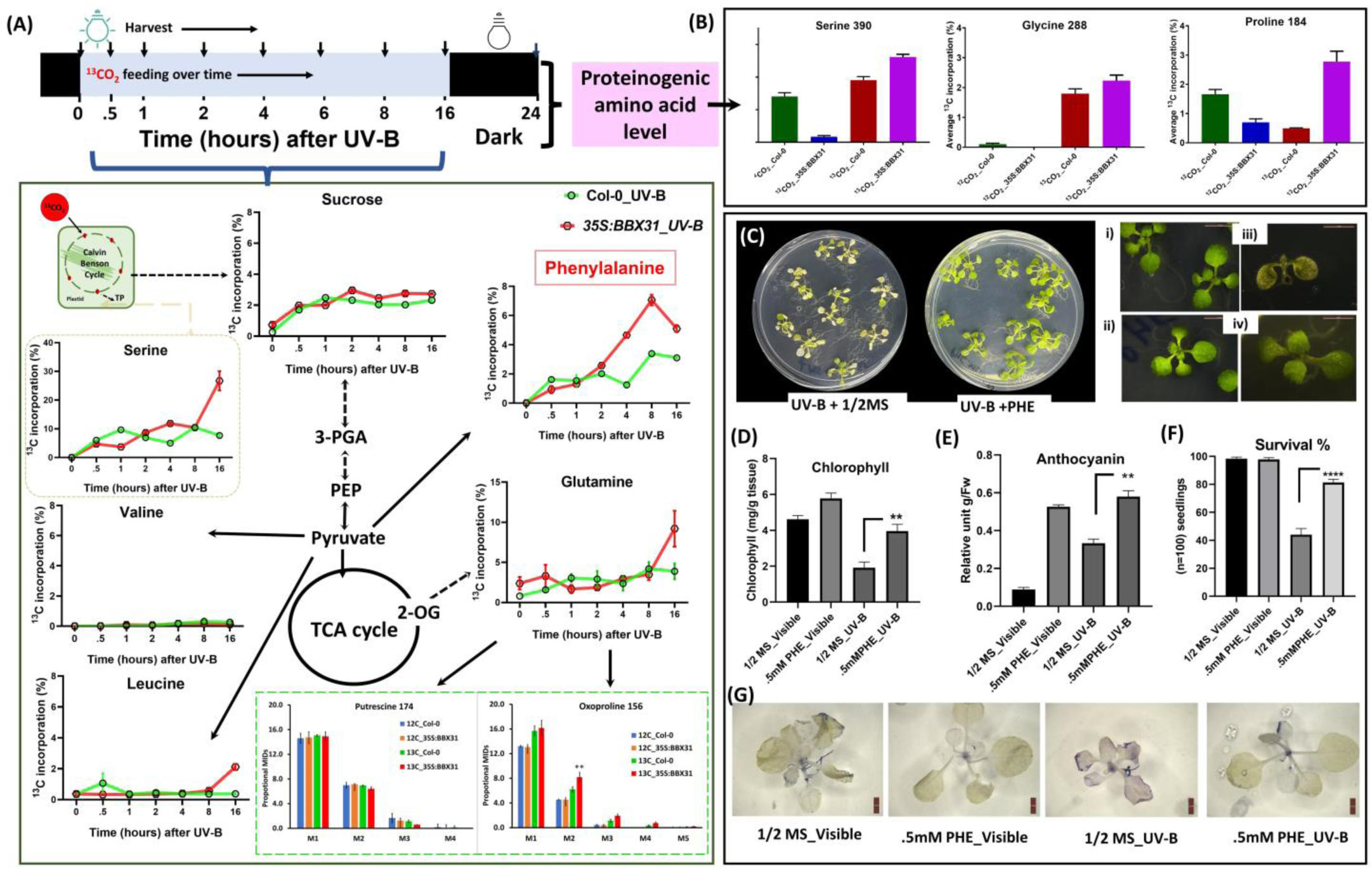
Kinetic ^13^CO_2_ metabolism in Arabidopsis Col-0 and BBX31 highlights denovo biosynthesis of key metabolites that confers tolerance under UV-B. *A)* The average 13C incorporation in the soluble metabolite’s fragments highlighted the higher activities of BBX31 in metabolism during UV-B treatment. In Col-0, the sucrose levels decreased more than *35S:BBX31* lines, demonstrating higher diminishing photosynthetic activity during UV-B. The 13C incorporation of glutamine and oxoproline was found to be higher in *35S:BBX31*. It has been observed that BCAA’s were not labeled under UV-B stress, even after long hours of 13C feeding. This suggests that the higher pools of BCAA’s might be coming from protein breakdown or other means, but not through denovo routes. The average 13C incorporation of oxoproline and putrescine was measured through mass isotopomer distributions of valid fragments, as shown in the green box. B)13C incorporation was depleted in proteinogenic amino acids as compared to visible, implying lower biosynthesis of protein biomass. However, levels of *35S:BBX31* proteinogenic amino acids were higher than Col-0. C) The 13C analysis also showed that the intracellular fate of carbon is routed towards the biosynthesis of phenylalanine during UV-B. Positive phenylalanine (PHE) effects were observed in growth parameters under UV-B light. When the Arabidopsis Col-0 plate was supplemented with phenylalanine, seedlings could survive in the plate when exposed to UV-B. The phenotype in the B panel is shown as i) Col-0 in MS plate, ii) supplemented with phenylalanine under visible light, iii) Col-0 in MS plate under UV-B, and iv) supplemented with phenylalanine under UV-B light. C) Under high UV-B treatment, phenylalanine rescued chlorophyll levels, E) upregulate higher anthocyanin levels and, F) resulted in a higher percentage of plants surviving. G) Cell death assay revealed that the phenylalanine supplementation can prevent tissue damage in Arabidopsis. This implies that carbon is directed towards the Phenyalaine to phenylpropanoid pathway and GS/GOGAT to produce proline, which helps BBX31 cope with stress under UV-B.

### BBX31 promotes accumulation of phenylalanine under UV-B

We integrated the data of metabolite levels, isotopomer distribution, and genotypes to analyse the phenylalanine levels under UV-B. Kinetic metabolomics showed cumulative trend of phenylalanine under UV-B exposure. The ^13^CO_2_ mapping studies also showed that the intracellular fate of carbon is routed towards biosynthesis of phenylalanine during UV-B stress and more incorporation of 13C was found in *35S:BBX31* **(Fig. 6A)**. Different experiments were designed to test the effects of phenylalanine (PHE) in susceptible Col-0 plants under UV-B **(Fig. 6C)**. Phenylalanine supplementing to Col-0 plants showed positive effects on growth parameters when exposed to UV-B. Phenylalanine rescued chlorophyll levels and upregulated higher anthocyanin levels **(Fig. 6D-F)**. Cell death assay revealed that the phenylalanine can prevent tissue damage in Arabidopsis **(Fig. 6G)**. Overall, it resulted in a higher percentage of surviving plants under UV-B.

## Discussion

UV-B can positively or negatively impact plant growth depending on the exposure time and dose. Determining photosynthetic assimilates is crucial to understanding plant phenotypes (Yadav *et al.,* 2020; Zeeman *et al.,* 2007). UV-B diminished photosynthetic compounds but BBX31 overexpressor lines have lesser chlorophyll degradation and retained more carotenoids compared to Col-0 and *bbx31* mutant. The abundant total phenolics and lowered IC50 value demonstrated the greater antioxidant activity of BBX31 under UV-B. Previous studies have shown that carotenoids and anthocyanins are the main pigments that protect plants from UV-B stress. They effectively scavenge reactive oxygen species and prevent photo-oxidative damage to chlorophyll caused by UV-B (Yamamoto 1996; Santin *et al.,* 2018; Lingwan *et al.,* 2023b). The role of B-box protein BBX31 in conferring UV-B tolerance stimulated us to investigate the metabolic readjustments in the tolerant lines of BBX31 compared to Col-0 upon exposure to UV-B. Comprehensive and dynamic metabolomics discriminated the metabolome among growth stages, and the variations in the levels of the metabolites triggered by UV-B conditions highlighted the potential reprogramming of central metabolic pathways in Arabidopsis.

### UV-B treatment reroutes the TCA cycle toward aspartate node

TCA intermediates are downregulated among all genotypes under UV-B stress irrespective of principle growth stage and genotypes. The relative levels of citrate, succinate, fumarate, malate was depleted and observed to be lowest in *35S:BBX31*. However, aspartate enhanced under early UV-B stress irrespective of genotypes. The metabolic levels of amino acids from the aspartate family pathway enhanced under UV-B treatments. The amino acids lysine, isoleucine, threonine levels were significantly abundant in the BBX31 genotype under UV-B. However, for defining the biosynthetic routes and carbon flux of Arabidopsis Col-0 and BBX31, It was essential to feed labelled CO_2_ under visible and UV-B. 13C analysis showed very low 13C incorporation under UV-B, suggesting TCA intermediates might be actively involved in carbon rerouting under UV-B stress. The TCA and Asp-family pathway plays a regulatory metabolic role in mitigating stress by maintaining cellular and energy metabolism. Previous studies suggest TCA decarboxylation was downregulated in highly illuminated leaves, wherein flux directed towards the enhanced biosynthesis of other essential amino acids (lysine, threonine and methionine) may be a strategy to maintain growth and stress tolerance (Tcherkez *et al.,* 2009; Atkin *et al.,* 2000; Tcherkez *et al.,* 2005; Galili, 2011). Our investigation suggested in the same direction that TCA intermediates are downregulated under UV-B. Aspartate family leads to the synthesis of the essential amino acids, whereas BBX31 highly enhances the levels of lysine, isoleucine, and threonine under UV-B tolerance and can open up new metabolic engineering possibilities towards the upregulation of essential amino acids in plants. However, in future, it will be interesting to see if the TCA cycle operates in the forward direction to form intermediated or if it functions differently under UV-B stress, e.g., it operates in both forward and reverse directions to produce glutamate via 2 oxoglutarate.

### BBX31 accumulates branched-chain amino acids to regulate cell metabolism under UV-B

Amino acid from the pyruvate pool-derived branched-chain amino acid (BCAAs) showed upregulation irrespective of growth stage under UV-B. The levels of valine and leucine were exceptionally abundant in *35S:BBX31*. In the metabolomics analysis, levels of lysine, isoleucine, threonine, alanine, leucine, and valine were enhanced under UV-B treatments. However, after ^13^CO_2_ labeling, we did not get any 13C incorporation in those amino acids. This suggested that the higher accumulation of those amino acids was via some catabolic routes to cope with UV-B stress, and there was no de novo carbon rerouted toward their biosynthesis **(Fig. 7A)**. In stress, oxidation of BCAAs plays a role of alternative source in reparation, supporting cellular integrity, and accumulation of valine plays a role in stress relief under UV-B. Certain reports have shown that amino acid catabolism is relevant whenever less amount of photosynthate source is available during stress in plants (Muñoz-Bertomeu *et al.,* 2013; Hildebrandt *et al.,* 2015; Igamberdiev and Kleczkowski, 2018). The oxidation of valine directly has a role as an electron donor into the electron transport chain (Binder, 2010). UV-B exposure alters the amino acid levels in photosynthetic organisms ranging from freshwater algae (Noaman, 2007), Barley (Valkama *et al.,* 2003), Rice (Kakani, 2003) and Arabidopsis (Kusano *et al.,* 2011).

**Fig. 7.**
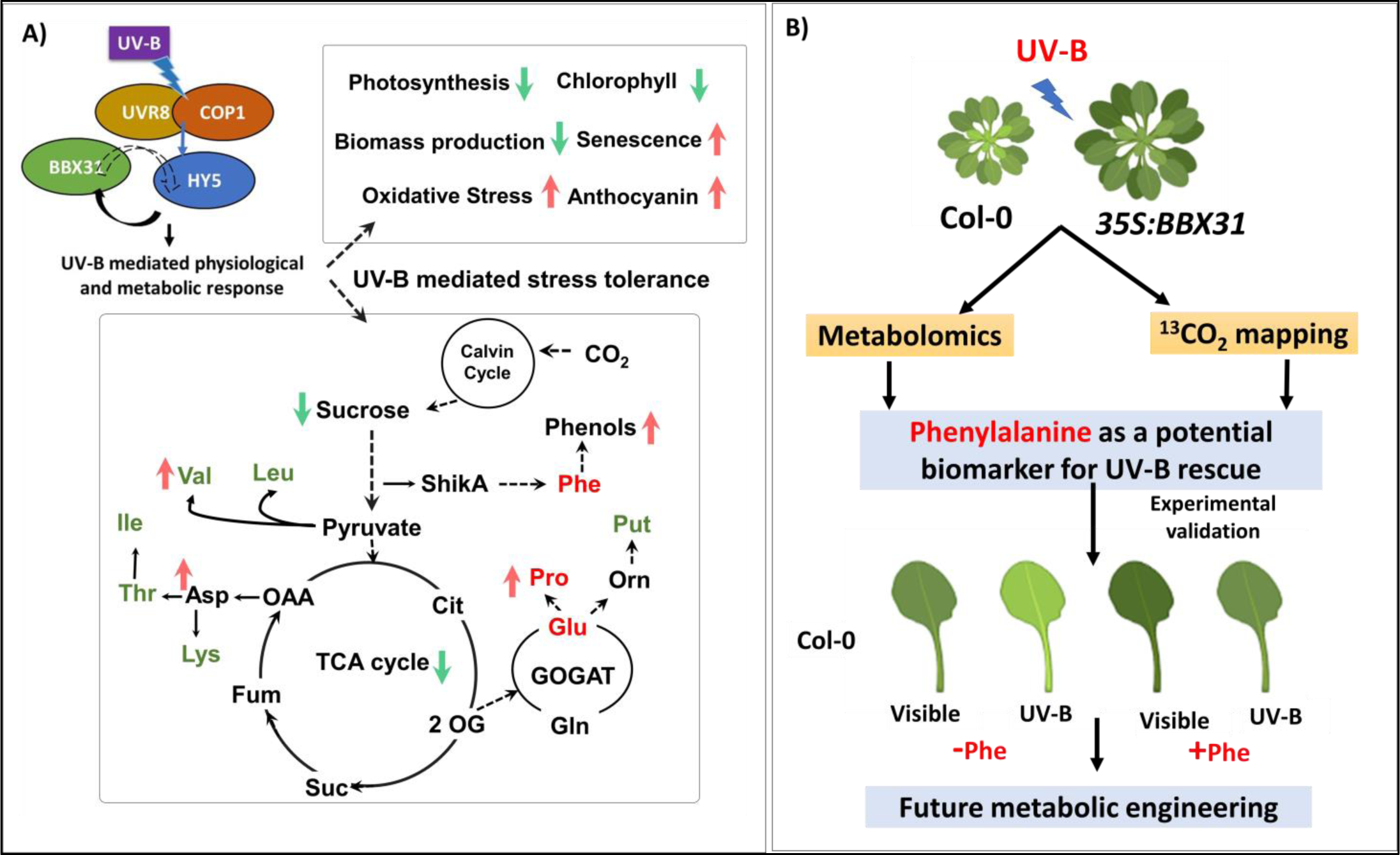
Metabolic reprogramming of UV-B conferred tolerance in Arabidopsis. A) On UV-B light, HY5 enhances BBX31 transcription by directly binding to its promoter. Reciprocally, overexpression of BBX31 enhances HY5 transcript levels in a UVR8-dependent manner. Black lines and dotted arrows indicate proven and possible transcriptional regulation. BBX31 influences the physiology and metabolic response under UV-B. Under UV-B exposure photosynthesis, chlorophyll levels, and biomass are reduced, while senescence, oxidative stress, anthocyanin, and other metabolic levels were enhanced. In comparison to the wild type, BBX31 overexpression promotes the accumulation of anthocyanin levels, amino acids, phenols, GS/GOGAT intermediates, and polyamines. ^13^CO_2_ feeding studies indicated there is no de novo biosynthesis of amino acids from pyruvate and aspartate node indicating remobilisation of pools of metabolic intermediates. Further, our studies confirmed BBX31 has higher 13C incorporation in phenylalanine, glutamine, and proline under UV-B, indicating rerouting of denovo carbon flux towards GS/GOGAT intermediates, phenylpropanoid, and shikimate pathways are known to play a crucial role in UV-B stress protection. Red upward arrows signify upregulation and green downward arrows represent downregulation of metabolic levels. Metabolites in green accumulated more from preexisting pools, while metabolites in red accumulation were rerouted from denovo biosynthesis. B) Integrating metabolomics, ^13^CO_2_ mapping studies showed denovo accumulation of phenylalanine, suggested to be a potential metabolic biomarker under UV-B. Phenylalanine (Phe) was found to have beneficial impacts on growth metrics when used under UV-B. These phenotypes suggested that Phe and metabolic pathways derived from Phe can be future targets for metabolic engineering toward UV-B tolerance crops. The illustration in Fig.7B was created using BioRender.

### BBX31 modulates polyamine metabolism to balance carbon-nitrogen

UV-B enhances GS/GOGAT metabolism, Glutamate plays a critical role in plant nitrogen, and central amino acid metabolism and acts as a substrate for glutamine biosynthesis. Glutamine biosynthesis may play a role in catabolism for further synthesis of polyamines in nitrogen balance and play a protective role in the seedlings (Forde and Lea, 2007; Limami *et al.,* 2008; Majumdar *et al.,* 2016; Kan *et. al.,* 2017). It forms a major network with ornithine, putrescine, oxoproline and other polyamines, which play a critical role in plant development and stress. In kinetic metabolic comparison to the wild type, the higher accumulation of proline, ornithine, citrulline and putrescine were observed in the *35S:BBX31*, showing their significant role under UV-B stress **(Fig. 7A)**. 13C labelling analysis showed significant 13C incorporation in glutamine and proline. Interestingly, we did not observe any 13C incorporation in putrescine. Therefore, the higher accumulation of putrescine under UV-B was not de novo biosynthesised. Previous studies in plants also highlight the protective role of glutamine accumulation under abiotic environmental stress conditions (May *et al.,* 1998; Paulose *et al.,* 2013). In UV-B stress, we observed oxoproline levels were modulated distinctly among Arabidopsis genotypes. The 13C analysis showed significant labeling in the oxoproline in BBX31 under UV-B. Given the oxo proline accumulation in Arabidopsis under UV-B and its metabolic link with glutathione (GSH) in plants, we speculate it is a key marker metabolite with a protective role in maintaining glutathione homeostasis. Glutathione homeostasis in plants provides stress tolerance (Paulose *et al.,* 2013; Khan *et al.,* 2019). In UV radiations, proline gets accumulated in the shoots of barley, rice, mustard, and mung bean seedlings (Saradhi *et al.,* 1995; Santin *et al.,* 2018). Overall, we observed that the 13C incorporation in glutamine, proline and oxoproline were high in *35S:BBX31*, implicating that it may have a role in mitigating UV-B stress response.

### UV-B exposure enhances photorespiration and modulates cell wall metabolism

Substantially high 13C label incorporation in glycine and serine indicates the possibility of active photorespiratory and glycolytic pathways in *35S:BBX31*. The pathway of serine formation initiates through a branch of glycolysis from 3-phosphoglycerate, and serine acts as a precursor for glycine formation. Accumulation of serine may play an important role during stress (Bykova *et al.,* 2005; Schaefer *et al.,* 1980). Soluble sugars play a vital role in stress tolerance in plants. The quantification of soluble sugars in Arabidopsis was, therefore, essential in understanding UV-B tolerance. Sucrose has low 13C enrichment, and in Col-0 levels was found to be decreased as compared to *35S:BBX31* lines; this shows low photosynthetic activity under UV-B. The levels of soluble sugars (fructose, glucose, erythrose, sucrose, myoinositol, glucopyranoside) are altered during UV-B stress. A subtle increment in some soluble sugars, which are involved in the composition of the cell wall matrix, is seen under UV-B in *35S:BBX31.* Similarly, enhanced levels of sugar alcohol in *35S:BBX31* such as sorbitol and erythritol may have a protective role under UV-B. Taken together, BBX31 in responses to UV-B modulates cell wall metabolism as defense plan to maintain cell wall integrity (Zhao *et al.,* 2016; Yadav *et al.,* 2019b).

### UV-B trigger phenylalanine accumulation and wax biosynthesis

The kinetic phenylalanine accumulation under UV-B may have direct effects on the biosynthesis or catabolism of phenylpropanoid intermediates. The 13C analysis also showed that the intracellular fate of carbon is routed towards the biosynthesis of de novo phenylalanine during UV-B stress. The highest incorporation of 13C was found in *35S: BBX31*. In addition, the phenylpropanoids derived metabolic pathway, the phenylalanine, sinapinic acid, benzoic acid, hydroxycinnamic acid, caffeic acid, vanillic acid, gallic acid, and coumaric acid respond strongly due to UV-B with the highest levels observed in *35S:BBX31* seedlings (Yadav *et al.,* 2019a). Phenylalanine account for almost ∼30% of the dry mass of a plant, In UV-B stress, enhancement in phenylalanine levels may be important to survival and playing a major role as a precursor to protein synthesis and phenylpropanoid mediates (Sullivan *et al.,* 2015). Pathway analysis provides insight that BBX31 also plays a role in the upregulation of cutin, suberin and wax biosynthesis. Previous studies have shown that many plants have developed features like cuticular wax and hairy structures to protect themselves from harmful UV-B (Saini *et al.,* 2020). All these results together indicate that UV-B has a major influence on the levels of secondary metabolites in plants. The elevated level of key phenolic secondary metabolite originating from the phenylpropanoid pathway is earlier reported to increase significantly in various medicinal plants (Kumari *et al.,* 2013; Zhang *et al.,* 2017; Taulavuori *et al.,* 2018; Lingwan *et al.,* 2023b)Taking together, capturing the early metabolic events in UV-B-mediated metabolite synthesis **(Fig. 7B)**, and accumulation can be practical for targeted phytochemical enrichment and for assisting metabolic engineering studies aimed at promoting stress tolerance in plants.

## Supplementary data

Additional supplementary information:

Fig. S1. Multivariate statistical analysis at the seedling stage of Arabidopsis Col-0, *bbx31* and *35S:BBX31* under visible and UV-B.

Fig.S2. Multivariate statistical analysis at the rosette growing stage of Arabidopsis Col-0, *bbx31* and *35S:BBX31* under visible and UV-B.

Fig. S3. Correlation analysis at the seedling and rosette stage

Fig.S4. Metabolic networks showed UV-B alters central metabolic pathways during the seedling stage.

Fig.S5. Metabolic levels of Col-0, *bbx31* mutant and *35S:BBX31* genotypes captured and visualized in active metabolic pathways during the rosette leaf growing stage.

Table S1. Metabolite Profiles during the seedling stage under Visible and UV-B.

Table S2. Metabolite Profiles during the rosette growing stage Visible and UV-B.

Table S3. Mapping the genes coding with enzymes revealed that BBX31 upregulates fatty acid biosynthesis.

Table S4. Joint pathway analysis visualised that BBX31 have a role in the upregulation of cutin, suberin and wax biosynthesis.

Table S5. Kinetic metabolite profiles captured during the rosette growing stage under visible light.

Table S6. Kinetic metabolite profiles captured during the rosette growing stage under UV-B treatment.

## Acknowledgements

SKM acknowledges the Science and Engineering Research Board (SERB) Early career research funding. SD would like to thank the Department of Biotechnology, Government of India, for funding. ML acknowledges the Indian Institute of Technology Mandi and the Government of India for a PhD fellowship, and AY acknowledges the UGC, Government of India, for a PhD fellowship.

## Author Contributions

ML and SKM conceptualized, developed methodology, analyzed data, and wrote the original draft. ML and AY conducted investigations. SKM and SD supervised the research. All authors contributed to writing, reviewing, and editing. SKM and SD secured funding.

## Data Availability

All supporting metadata is available in the manuscript figures and supplementary materials.

## References

Allen DK, Libourel IG, Shachar-Hill YA. 2009. Metabolic flux analysis in plants: coping with complexity. Plant, cell & environment, 329.:1241–57.

Arnon AN. 1967. Method of extraction of chlorophyll in the plants. Agronomy journal 23.1: 112–121.

Binder S. 2010. Branched-Chain Amino Acid Metabolism in Arabidopsis thaliana. The arabidopsis book 8: e0137-e0137

Boyes DC, Zayed AM, Ascenzi R, McCaskill AJ, Hoffman NE, Davis KR, Görlach J. 2001. Growth stage-based phenotypic analysis of Arabidopsis: a model for high throughput functional genomics in plants. The Plant cell 13: 1499–1510

Bykova NV, Keerberg O, Pärnik T, Bauwe H, Gardeström P. 2005. Interaction between photorespiration and respiration in transgenic potato plants with antisense reduction in glycine decarboxylase. Planta 222: 130–140

Caldwell MM, Robberecht R, Flint SD. 1983. Internal filters: Prospects for UV-acclimation in higher plants. Physiologia Plantarum 58: 445–450

Caldwell MM, Teramura AH, Tevini M. 1989. The changing solar ultraviolet climate and the ecological consequences for higher plants. Trends in Ecology & Evolution 4:363–367

Celeste Dias M, Pinto DCGA, Correia C, Moutinho-Pereira J, Oliveira H, Freitas H, Silva AMS, Santos C. 2018. UV-B radiation modulates physiology and lipophilic metabolite profile in Olea europaea. Journal of Plant Physiology 222: 39–50

Chun OK, Kim DO, Lee CY. 2003. Superoxide radical scavenging activity of the major polyphenols in fresh plums. Journal of agricultural and food chemistry, 5127., 8067–8072.

Chuwah C, van Noije T, van Vuuren DP, Stehfest E, Hazeleger W. 2015. Global impacts of surface ozone changes on crop yields and land use. Atmospheric Environment 106: 11–23

Coohill TP. 1989. Ultraviolet Action Spectra 280 To 380 Nm. And Solar Effectiveness Spectra for Higher Plants. Photochemistry and Photobiology 50: 451–457

Forde BG, Lea PJ. 2007. Glutamate in plants: metabolism, regulation, and signalling. Journal of Experimental Botany 58: 2339–2358

Galili G. 2011. The aspartate-family pathway of plants: linking production of essential amino acids with energy and stress regulation. Plant signaling & behavior 6: 192–195

Gangappa SN, Botto JF. 2014. The BBX family of plant transcription factors. Trends Plant Science 19: 460–470

Gutbrod K, Romer J, Dörmann P. 2019. Phytol metabolism in plants. Progress in lipid research, 74: 1–17

Ha J, Park JY, Choi Y, Chang PS, Park KM. 2021. Comparative Analysis of Universal Protein Extraction Methodologies for Screening of Lipase Activity from Agricultural Products. Catalysts, 117., 816.

Hasunuma T, Harada K, Miyazawa SI, Kondo A, Fukusaki E, Miyake C. 2010. Metabolic turnover analysis by a combination of in vivo 13C-labelling from ^13^CO_2_ and metabolic profiling with CE-MS/MS reveals rate-limiting steps of the C3 photosynthetic pathway in Nicotiana tabacum leaves. Journal of Experimental Botany, 614., 1041–1051.

Hildebrandt Tatjana M, Nunes Nesi A, Araújo Wagner L, Braun HP. 2015. Amino Acid Catabolism in Plants. Molecular Plant 8: 1563–1579

Igamberdiev AU, Kleczkowski LA. 2018. Corrigendum: The Glycerate and Phosphorylated Pathways of Serine Synthesis in Plants: The Branches of Plant Glycolysis Linking Carbon and Nitrogen Metabolism. Frontiers in Plant Science, 9

Jansen MAK, Gaba V, Greenberg BM. 1998. Higher plants and UV-B radiation: balancing damage, repair and acclimation. Trends in Plant Science 3: 131–135

Jenkins GI. 2017. Photomorphogenic responses to ultraviolet-B light. Plant Cell Environ 40: 2544–2557

Job N, Lingwan M, Masakapalli SK, Datta S. 2022. Transcription factors BBX11 and HY5 interdependently regulate the molecular and metabolic responses to UV-B. Plant Physiology. 1;189(4):2467–80.

Kakani VG 2003. Field crop responses to ultraviolet-B radiation: a review. Agricultural and forest meteorology v. 120: pp. 191-218-2003 v.2120 no.2001-2004

Kan CC, Chung TY, Wu HY, Juo YA, Hsieh MH. 2017. Exogenous glutamate rapidly induces the expression of genes involved in metabolism and defense responses in rice roots. BMC genomics 18: 186–186

Kataria S, Jajoo A, Guruprasad KN. 2014. Impact of increasing Ultraviolet-B UV-B. radiation on photosynthetic processes. Journal of Photochemistry and Photobiology B: Biology 137: 55–66

Khan N, Bano A, Babar MA. 2019. Metabolic and physiological changes induced by plant growth regulators and plant growth promoting rhizobacteria and their impact on drought tolerance in *Cicer arietinum* L. PloS one 14

Khanna R, Kronmiller B, Maszle DR, Coupland G, Holm M, Mizuno T, Wu SH. 2009. The Arabidopsis B-box zinc finger family. Plant Cell, 2111.:3416–20.

Kliebenstein DJ, Lim JE, Landry LG, Last RL 2002. Arabidopsis UVR8 regulates ultraviolet-B signal transduction and tolerance and contains sequence similarity to human regulator of chromatin condensation. Plant Physiology, 130: 234–243

Koley S, Chu KL, Mukherjee T et al. 2022. Metabolic synergy in Camelina reproductive tissues for seed development. Science advances. 28;8(43):eabo7683.

Kopka J, Schauer N, Steinhauser D et al. 2004.: the Golm Metabolome Database. Bioinformatics, 21: 1635–1638

Kumari A, Parida AK. 2018. Metabolomics and network analysis reveal the potential metabolites and biological pathways involved in salinity tolerance of the halophyte Salvadora persica. Environmental and Experimental Botany, 148: 85–99

Kumari R, Prasad MN. 2013. Medicinal plant active compounds produced by UV-B exposure. Sustainable Agriculture Reviews: Volume 12; 225–54.

Kupina S, Fields C, Roman MC, Brunelle SL. 2019. Determination of total phenolic content using the Folin-C assay: Single-laboratory validation. Journal of AOAC International 1015., 1466–1472.

Kusano M, Tohge T, Fukushima A, Fernie AR et al. 2011. Metabolomics reveals comprehensive reprogramming involving two independent metabolic responses of Arabidopsis to UV-B light. The Plant Journal, 67: 354–369

Kushwaha AK, Dwivedi S, Mukherjee A, Lingwan M, Dar MA, Bhagavatula L, Datta S. 2022. Plant microProteins: Small but powerful modulators of plant development. Iscience. 18;25(11).

Limami AM, Glévarec G, Ricoult C, Cliquet J-B, Planchet E. 2008. Concerted modulation of alanine and glutamate metabolism in young Medicago truncatula seedlings under hypoxic stress. Journal of experimental botany, 59: 2325–2335

Lingwan M, Shagun S, Masakapalli SK et al. 2023. Phytochemical rich Himalayan Rhododendron arboreum petals inhibit SARS-CoV-2 infection in vitro. Journal of Biomolecular Structure and Dynamics, 4;41(4):1403–13.

Lingwan M, Masakapalli SK. 2022. A robust method of extraction and GC-MS analysis of Monophenols exhibited UV-B mediated accumulation in Arabidopsis. Physiology and Molecular Biology of Plants. 28(2):533–43.

Lingwan M, Pradhan AA, Kushwaha AK, Dar MA, Bhagavatula L, Datta S. 2023. Photoprotective role of plant secondary metabolites: Biosynthesis, photoregulation, and prospects of metabolic engineering for enhanced protection under excessive light. Environmental and Experimental Botany, 209, 105300.

Lingwan, M. (2024). Damage and Repair: How Poaceae plants fix DNA damaged by UV-B radiation. Plant physiology, 10.1093/plphys/kiae094

Lisec J, Schauer N, Kopka J, Willmitzer L, Fernie AR. 2006. Gas chromatography mass spectrometry–based metabolite profiling in plants. Nature Protocols 1: 387–396

Lommen A, Kools HJ. 2012. MetAlign 3.0: performance enhancement by efficient use of advances in computer hardware. Metabolomics 8: 719–726

Luo W, Brouwer C. 2013. Pathview: an R/Bioconductor package for pathway-based data integration and visualization. Bioinformatics 29: 1830–1831

Ma F, Jazmin LJ, Young JD, Allen DK. 2017. Isotopically nonstationary metabolic flux analysis INST-MFA. of photosynthesis and photorespiration in plants. In Photorespiration pp. 167-194. Humana Press, New York, NY.

Maiti S, Moon UR, Bera P, Samanta T, Mitra A. 2014. The in vitro antioxidant capacities of Polianthes tuberosa L. flower extracts. Acta Physiologiae Plantarum 36, 2597–2605.

Majumdar R, Barchi B, Turlapati SA, Gagne M, Minocha R, Long S, Minocha SC. 2016. Glutamate, Ornithine, Arginine, Proline, and Polyamine Metabolic Interactions: The Pathway Is Regulated at the Post-Transcriptional Level. Frontiers in Plant Science, 7

Masakapalli SK, Bryant FM, Kruger NJ, Ratcliffe RG. 2014. The metabolic flux phenotype of heterotrophic Arabidopsis cells reveals a flexible balance between the cytosolic and plastidic contributions to carbohydrate oxidation in response to phosphate limitation. The Plant Journal, 78: 964–977

Masakapalli SK, Kruger NJ, Ratcliffe RG. 2013. The metabolic flux phenotype of heterotrophic Arabidopsis cells reveals a complex response to changes in nitrogen supply. The Plant Journal, 744., 569–582, 2013.

Masakapalli SK. 2011. Network flux analysis of central metabolism in plants Doctoral dissertation, Oxford University, UK.

May MJ, Vernoux T, Leaver C, Montagu MV, Inzé D. 1998. Glutathione homeostasis in plants: implications for environmental sensing and plant development. Journal of Experimental Botany, 49: 649–667

Munoz-Bertomeu J, Anoman A, Flores-Tornero M, Toujani W, Rosa-Téllez S, Fernie AR, Roje S, Segura J, Ros R. 2013. The essential role of the phosphorylated pathway of serine biosynthesis in Arabidopsis. Plant signaling & behavior, 8: e27104–e27104

Noaman N. 2007. Ultraviolet-B Irradiation Alters Amino Acids, Proteins, Fatty Acids Contents and Enzyme Activities of Synechococcus leopoliensis. International Journal of Botany

Pang Z, Chong J, Zhou G, de Lima Morais, DA, Chang L, Barrette M, Xia J. 2021. MetaboAnalyst 5.0: narrowing the gap between raw spectra and functional insights. Nucleic acids research, 49(W1), W388–W396

Pant Y, Lingwan M, Masakapalli SK. 2023. Metabolic, biochemical, mineral and fatty acid profiles of edible Brassicaceae microgreens establish them as promising functional food. Food Chemistry Advances, 3, 100461.

Parihar P, Singh S, Singh R, Singh VP, Prasad SM. 2015. Changing scenario in plant UV-B research: UV-B from a generic stressor to a specific regulator. Journal of photochemistry and photobiology. B, Biology 153: 334–343

Paulose B, Chhikara S, Coomey J, Jung HI, Vatamaniuk O, Dhankher OP. 2013. A γ-glutamyl cyclotransferase protects Arabidopsis plants from heavy metal toxicity by recycling glutamate to maintain glutathione homeostasis. The Plant Cell, 1;25(11):4580–95.

Rai N, O’Hara A, Farkas D, et al. 2020. The photoreceptor UVR8 mediates the perception of both UV-B and UV-A wavelengths up to 350 nm of sunlight with responsivity moderated by cryptochromes. Plant, cell & environment. 43(6):1513–27.

Reis-Mansur M, Cardoso-Rurr JS. 2019. Carotenoids from UV-resistant Antarctic Microbacterium sp. LEMMJ01. Scientific reports, 9(1), 9554

Reumann S, Weber APM. 2006. Plant peroxisomes respire in the light: Some gaps of the photorespiratory C2 cycle have become filled—Others remain. Biochimica et Biophysica Acta BBA. - Molecular Cell Research, 1763: 1496–1510

Robson TM, Klem K, Urban O, Jansen MAK. 2015. Re-interpreting plant morphological responses to UV-B radiation. Plant, Cell & Environment 38: 856–866

Saini P, Bhatia S, Mahajan M, Kaushik A. et al. 2020. ELONGATED HYPOCOTYL5 negatively regulates DECREASE WAX BIOSYNTHESIS to increase survival during UV-B stress. Plant physiology. 1;184(4):2091–106.

Sakamoto M, & Suzuki T. 2017. Synergistic effects of a night temperature shift and methyl jasmonate on the production of anthocyanin in red leaf lettuce. American Journal of Plant Sciences, 807., 1534.

Schaefer J, Kier LD, Stejskal EO. 1980. Characterization of photorespiration in intact leaves using 13carbon dioxide labeling. Plant physiology, 652., 254–259.

Schwender J, 2008. Metabolic flux analysis as a tool in metabolic engineering of plants. Current Opinion in Biotechnology, 192., pp.131–137.

Shree M, Masakapalli SK. 2018. Intracellular fate of universally labelled 13C isotopic tracers of glucose and xylose in central metabolic pathways of Xanthomonas oryzae. Metabolites. 15;8(4):66.

Shree M, Lingwan M, Masakapalli SK. 2019. Metabolite profiling and metabolomics of plant systems using 1H NMR and GC-MS. OMICS-Based Approaches in Plant Biotechnology, 129, 129.

Song Z, Yan T, Liu J, Bian Y, Heng Y, Lin F, Jiang Y, Wang Deng X, Xu D. 2020. BBX28/BBX29, HY5 and BBX30/31 form a feedback loop to fine-tune photomorphogenic development. The Plant Journal, 104: 377–390

Sullivan JH, Muhammad D, Warpeha KM. 2015. Phenylalanine Is Required to Promote Specific Developmental Responses and Prevents Cellular Damage in Response to Ultraviolet Light in Soybean Glycine max. during the Seed-to-Seedling Transition. PLOS ONE, 9: e112301

Taulavuori K, Pyysalo A, Taulavuori E, Julkunen-Tiitto R. 2018. Responses of phenolic acid and flavonoid synthesis to blue and blue-violet light depends on plant species. Environmental and Experimental Botany 150: 183–187

Tcherkez G, Cornic G, Bligny R, Gout E, Ghashghaie J. 2005. In vivo respiratory metabolism of illuminated leaves. Plant Physiol 138: 1596–1606

Tcherkez G, Mahé A, Gauthier P, Mauve C, Gout E, Bligny R, Hodges M. 2009.. In folio respiratory fluxomics revealed by 13C isotopic labeling and H/D isotope effects highlight the noncyclic nature of the tricarboxylic acid “cycle” in illuminated leaves. Plant Physiology, 1512., 620–630.

Vaishak KP, Yadukrishnan P, Bakshi S, Kushwaha AK, Ramachandran H, Job N, Babu D, Datta S. 2019. The B-box bridge between light and hormones in plants. Journal of Photochemistry and Photobiology B: Biology 191: 164–174

Valkama E, Kivimäenpää M, Hartikainen H, Wulff A. 2003. The combined effects of enhanced UV-B radiation and selenium on growth, chlorophyll fluorescence and ultrastructure in strawberry Fragaria x ananassa. and barley Hordeum vulgare. treated in the field. Agricultural and forest meteorology. 120: 267–278

Yadav A, Bakshi S, Yadukrishnan P, Lingwan M, Dolde U, Wenkel S, Masakapalli SK, Datta S. 2019. The B-Box-Containing MicroProtein miP1a/BBX31 Regulates Photomorphogenesis and UV-B Protection. Plant Physiology 179: 1876

Yadav A, Lingwan M, Yadukrishnan P, Masakapalli SK, Datta S. 2019. BBX31 promotes hypocotyl growth, primary root elongation and UV-B tolerance in Arabidopsis. Plant Signaling & Behavior 14: e1588672

Yadav A, Singh D, Lingwan M, Yadukrishnan P, Masakapalli SK, Datta, S. 2020. Light signaling and UV-B-mediated plant growth regulation. Journal of integrative plant biology, 629., 1270-1292.

Yoshiyama KO, Sakaguchi K, Kimura S. 2013. DNA damage response in plants: conserved and variable response compared to animals. Biology 2: 1338–1356

Zeeman SC, Smith SM, Smith AM. 2007. The diurnal metabolism of leaf starch. Biochemical Journal, 4011.:13–28.

Zhang X, Huai J, Shang F, Xu G, Tang W, Jing Y, Lin R. 2017. A PIF1/PIF3-HY5-BBX23 Transcription Factor Cascade Affects Photomorphogenesis. Plant Physiology, 174: 2487–2500

Zhao L, Chanon AM, Chattopadhyay N, Dami IE, Blakeslee JJ. 2016. Quantification of Carbohydrates in Grape Tissues Using Capillary Zone Electrophoresis. Frontiers in plant science 7: 818–818

